# Relevance of the iron-responsive element (IRE) pseudotriloop structure for IRP1/2 binding and validation of IRE-like structures using the yeast three-hybrid system

**DOI:** 10.1101/524991

**Authors:** Shih-Cheng Chen, René C.L. Olsthoorn

## Abstract

Iron-responsive-elements (IREs) are ~35-nucleotide (nt) stem-loop RNA structures located in 5′ or 3′ untranslated regions (UTRs) of mRNAs, and mediate post-transcriptional regulation by their association with IRE-binding proteins (IRPs). IREs are characterized by their apical 6-nt loop motif 5′-CAGWGH-3′ (W = A or U and H= A, C or U), the so-called pseudotriloop, of which the loop nts C1 and G5 are paired, and the none-paired C between the two stem regions. In this study, the yeast three-hybrid (Y3H) system was used to investigate the relevance of the pseudotriloop structure of ferritin light chain (FTL) for the IRE-IRP interaction and the binding affinities between variant IRE(-like) structures and the two IRP isoforms, IRP1 and 2. Mutational analysis of FTL IRE showed that deletion of the bulged-out U6 of the pseudotriloop does not significantly affect its binding to either IRP1 or 2, but substitution with C enhances binding of both IRPs. In addition, IRP1 was found more sensitive toward changes in the pseudotriloop-stabilizing C1-G5 base pair than IRP2, while mutation of the conserved G3 was lowering the binding of both IRPs. In comparison to FTL IRE other variant IREs, IRE of 5′-aminolevulinate synthase 2 (ALAS2), SLC40A1 (also known as Ferroportin-1), and endothelial PAS domain protein 1 (EPAS1) mRNA showed slightly higher, similar, and slightly weaker affinity for IRPs, respectively, while SLC11A2 IRE exhibited very weak binding to IRP1 and medium binding to IRP2, indicating the different binding modes of IRP1 and 2. Notably, α-Synuclein IRE showed no detectable binding to either IRP1 or 2. Our results indicate that Y3H represents a *bona fide* system to characterize binding between IRPs and various IRE-like structures.

## Introduction

Iron (Fe^2+^) is involved in multiple cellular processes, including respiration, DNA synthesis, oxygen transport, energy metabolism, *etc.*, and hence its cellular homeostasis is precisely regulated in most organisms. In vertebrates, the cellular iron level is stably maintained within a narrow range to avoid damage to protein, DNA, and lipid, as an excess or deficiency of cellular iron may lead to physiological misregulation, *e.g*. abnormal cell proliferation. In humans, impaired iron homeostasis is associated with inherited hemochromatosis, cirrhosis, cardiomyopathy, diabetes mellitus, and many neurodegenerative disorders such as Friedreich’s ataxia, Parkinson’s and Alzheimer’s diseases [1–3]. The uptake of cellular iron is predominantly accomplished by divalent metal transporter 1 (DMT1, also known as SLC11A2 or NRAMP2) and transferrin receptor (TfR1), while the storage and sequestration of cellular iron is mediated by H- and L-ferritin subunits (HTL and FTL), and the export and translocation of cellular iron is facilitated by the basolateral exporter Ferroportin (encoded by SLC40A1). All these actions are tightly controlled to regulate cellular iron homeostasis (reviewed in [4]).

The key regulator of intracellular iron metabolism is Iron-Regulatory Protein (IRP), a member of the aconitase superfamily that post-transcriptionally regulates many genes involved in iron metabolism by specifically binding to the conserved Iron Responsive Elements (IREs) located in the UTRs of mRNAs [5–7]. For instance, when Fe^2+^ levels are low, IRPs are able to tightly bind IRE(s) present in the 3′ UTR of the TfR1 mRNA to stabilize the transcripts and increase the protein level of TfR1 that promotes iron uptake. On the other hand, low Fe^2+^ levels also facilitate IRP to interact with IREs located in the 5′ UTR of SLC40A1 mRNA and reduce the protein level of the iron-exporting Ferroportin. As a result, the iron-sensing IRP-IRE interaction increases the uptake and the availability of iron in cells. On the contrary, high Fe^2+^ levels decrease IRP’s IRE-binding affinity, resulting in reduced stability of TfR1 mRNA and increased translation of SLC40A1 mRNA that synergistically depletes cellular iron [8]. Thus, knowledge of IRE binding affinity is key to the study of IRP-mediated iron homeostasis [9].

The sequence and structure of IREs are highly conserved in mammalians, and generally consist of a stem-loop element formed by a five base pair stem carrying a 6-nucleotide (nt) apical loop motif 5′-CAGWGH-3′ (W = A or U and H= A, C or U) separated from a lower stem of variable length by an internal loop or bulge containing a conserved C [6, 10–12]. Since the classic IREs have been identified in ferritin mRNA[13, 14], quite a few IRE-like structures have been proposed later in other mRNAs, of which the encoded proteins are not necessarily solely involved in iron homeostasis, *e.g*. 5′-aminolevulinate synthase 2 (ALAS2), endothelial PAS domain protein 1 (EPAS1), and α-Synuclein [15–19]. Some of these RNA elements, discovered by immunoprecipitation, were demonstrated to have IRP binding affinity *in vitro* using electrophoretic mobility shift assays (EMSA), however, those predicted by computational analysis, like the IRE-like structure found in α-Synuclein mRNA, have not been reported to bind IRPs.

In this study, we took advantage of the yeast three-hybrid (Y3H) system [20, 21] which has been broadly used to assay RNA-protein interaction *in vivo* [22, 23] to study the relevance of the pseudotriloop conformation of the FTL IRE for binding of IRP1 and 2. Furthermore, we investigated the binding potential of several IRE-like structures to both IRP isoforms.

## Materials and Methods

### Constructs of RNA and protein expressing vectors for yeast three-hybrid system

The Y3H system used in this study is generally adapted from what has been reported by SenGupta *et al.*, (1996). The wildtype FTL (FTL-wt) IRE is taken as a standard IRE in this study, of which the apical loop consists of 5′-CAGUGU-3′. IRE mutants and variants were constructed by cloning complementary pairs of chemically synthesized DNA oligonucleotides (Eurogentec, Belgium) into NheI/BglII digested pIIIA/MS2.1*, a modified version of pIIIA/MS2.1 [24] in which the unique SmaI and the flanking redundant sequences were removed by replacing the fragment between the two EcoRI sites with the sequence 5′-aatttatactcacatgaggatcacccatgtaattaacactgaggatcacccagtggctagcttctagaaagatctg-3′ thereby creating unique NheI, XbaI, and BglII sites downstream of the MS2 CP binding sequences [25]. Additional nucleotides were introduced at the bottom of the lower stem of the IREs (indicated in Fig. 2 and 6) that stabilize the RNA structure in order to minimize the potential formation of alternative structures. The RNA-binding protein expressing vector was based on the shuttle vector pGADT7 (Clontech, USA) and constructed as described previously [25]. DNA fragments encoding IRP1 and 2 were generated by PCR and inserted between SmaI and XhoI sites of pGADT7, in frame with the GAL4 domain at the C-terminus.

### Yeast three-hybrid assay

The pIIIA/MS2.1-IREs, vectors that produce the chimeric RNA of IRE and MS2 Coat protein (CP) binding sequence, and the pGADT7-IRPs, vectors that express the IRP-GAL4 fusion protein were co-introduced to yeast strain L40-coat which expresses the MS2-CP-LexA fusion protein [21]. Transformed yeast cells inheriting both vectors were cultured on selective medium. To assay the reporter of RNA-protein interaction, HIS3 expression, we monitored the phenotypic growth of transformed yeast L40 coat on 3-Amino-1,2,4-triazole (3-AT) containing Yeast Nitrogen Base (YNB) medium as described previously [25, 26]. Briefly, three independent colonies of L40-coat yeast which were transformed simultaneously with pIIIA/MS2.1-IREs and pGADT7-IRPs were cultured 2 days in Leu^−^, Ade^−^, and − Ura^−^ YNB medium at 30°C. The yeasts were then sub-cultured, in 1-to-1,000 ratio, to an optical density of ~0.2 at 600 nm, followed by applying 2 μl onto Leu^−^, Ade^−^, Ura^−^, and His^−^ YNB agar containing 0 to 15 mM 3-AT. After culturing for 4 days at 30°C, the phenotypic growth of two represented clones was documented.

## Results

### Binding affinity between IREs and IRPs in yeast three-hybrid system

The RNA baits, FTL-wt IRE and its mutants, and the protein preys, IRP1 and 2, were cloned into expression vectors for Y3H assay, respectively (Fig. 1, adapted from SenGupta *et al.*[21]). Each mutant of FTL IRE (Fig. 2) was co-transformed to yeast strain L40-Coat with either IRP1 or IRP2 expressing vectors to study the IRE-IRP interaction in yeast. The binding affinity between the tested IREs and IRPs was evaluated by the phenotypic growth of the yeast on HIS^−^ media, as binding between the bait and the prey leads to trans-activation of the reporter gene *HIS3* [21, 23, 24], enabling the transformed yeast to grow on media without histidine supplementation.

**Figure 1.**
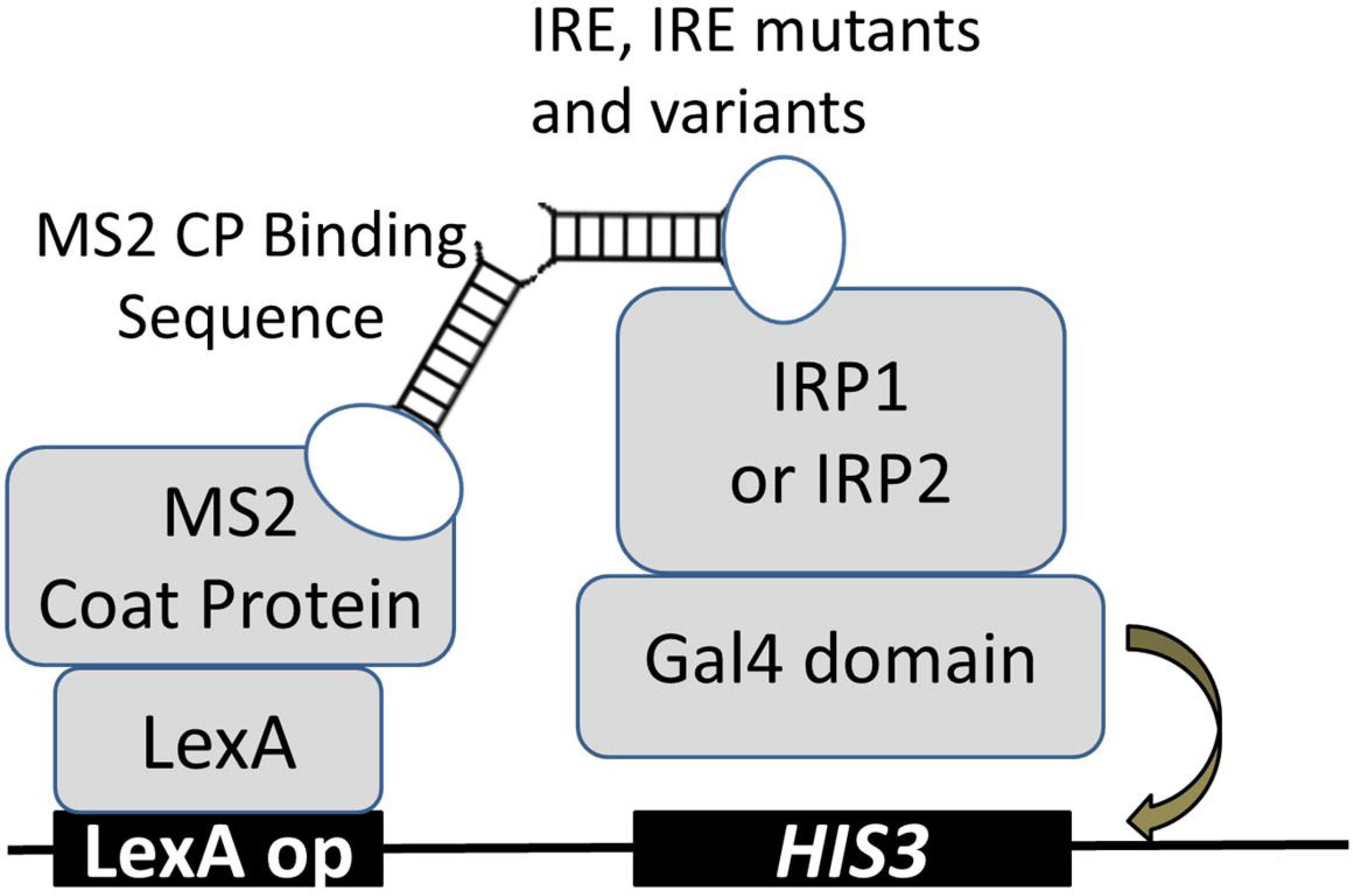
The yeast three-hybrid (Y3H) system to assay binding affinity between IRE and IRP1 and 2. The principle components of the Y3H system used in this study are shown, including (i) the hook, fusion protein consisting of LexA DNA-binding domain and MS2 coat protein (CP), (ii) the bait, a chimeric RNA structure consists of MS2 CP binding sequence and IRE, IRE mutants or variants, (iii) the fish, transcriptional activation domain Gal4 fused to IRP1 or 2. The transcriptional activation of the reporter gene HIS3 only occurs upon sufficient RNA-protein interaction between the tested IRE and the IRP that completes the formation of the trimeric complex (illustration adapted form [21]).

**Figure 2.**
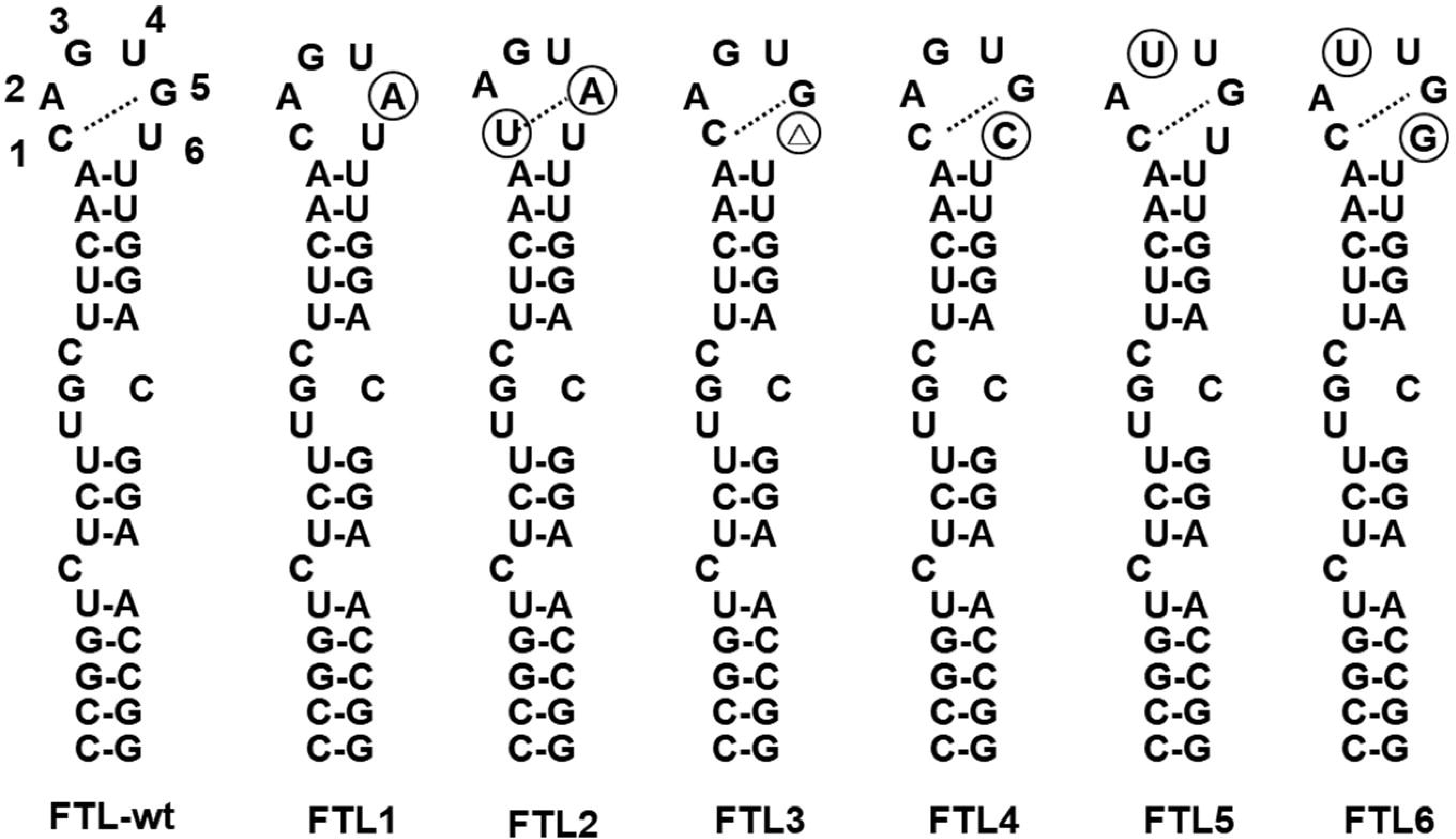
The IRE mutants tested in this study. The secondary structures of the IRE of ferritin light chain (FTL-wt) and the mutants tested in this study (FTL1-6) are shown. Base substitutions are marked with a circle while base deletions are marked with a triangle. The dotted-line represents the intra-loop base pairing of the pseudotriloop.

In Y3H assays, non-specific RNA-protein/protein-protein interaction or promoter leakage may lead to background HIS3 expression. Therefore, 3-AT is usually added to the medium to prevent yeasts from growing independently of a positive RNA-protein interaction. In our study of IRE-IRP interaction, yeast possessing solely MS2-coat binding sequence without the RNA bait IRE, showed false positive growth on selective medium without 3-AT when the vector expressing either IRP1 or 2 was co-transformed (Fig. 3). However, addition of 2 mM of 3-AT could completely suppress such false positive growth and can be considered as the threshold for specific IRE-IRP interactions in the following tests. IRP1 shows strong binding affinity to FTL-wt IRE, exhibiting good growth on the media containing 3-AT up to 6 mM (Fig. 3). A similar growth pattern was observed for IRP2 (Fig. 3), suggesting that IRP2 and IRP1 have similar binding affinity for the wildtype FTL IRE.

**Figure 3.**
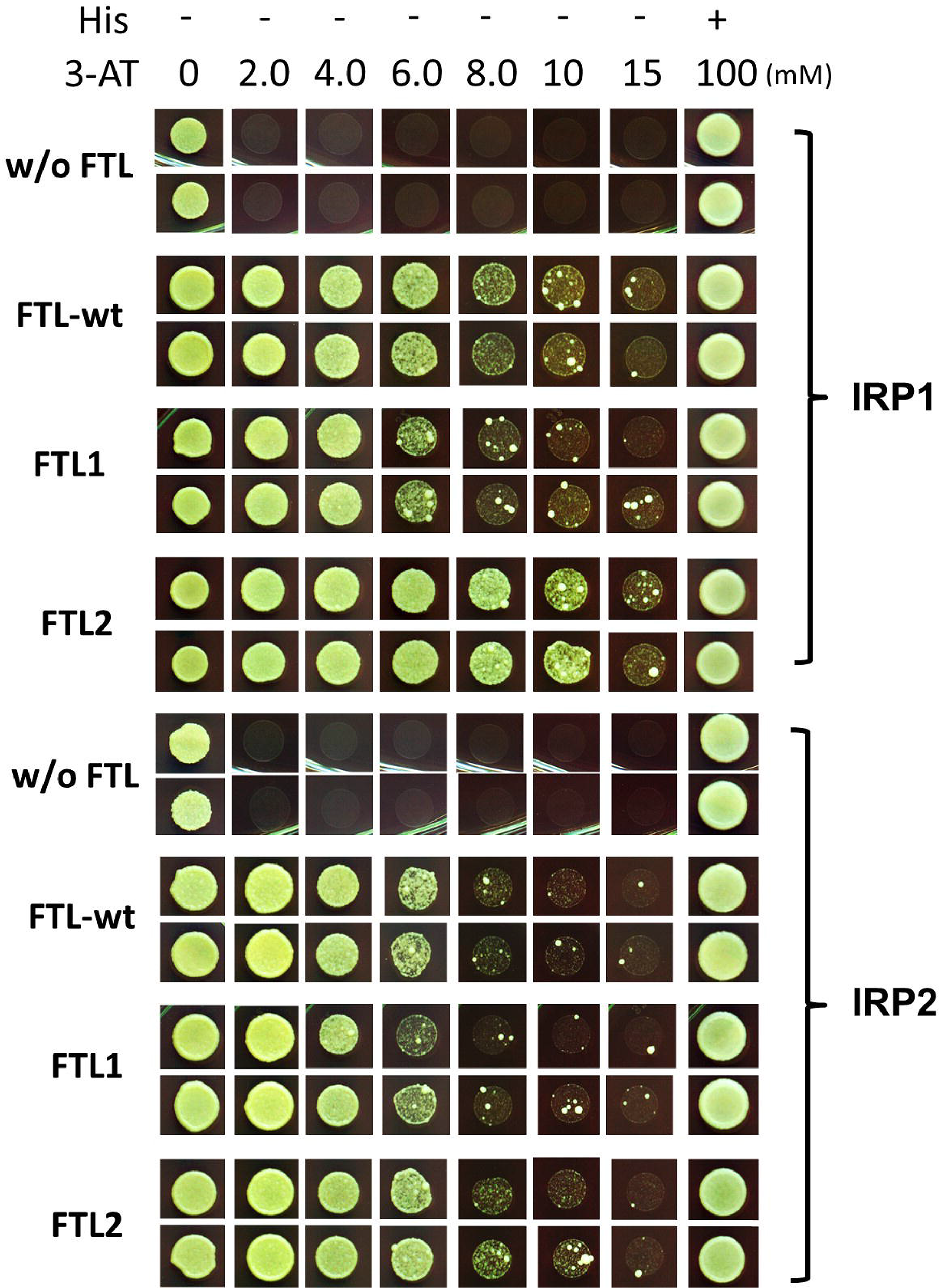
Yeast three-hybrid assay to verify the relevance of the C1-G5 base pair of the IRE pseudotriloop to the binding with IRP1 and 2. In the yeast assay, higher resistance to 3-AT represents a stronger RNA-protein interaction. “w/o FTL” is the negative control of which the RNA ligand has no IRE structure, while clones FTL1 and FTL2 contain IRE mutants shown in Fig. 2. The upper panel displays the binding affinities between the tested FTL IRE/mutants and IRP1 while the lower panel shows the binding to IRP2, respectively.

### Relevance of the IRE pseudotriloop structure for IRP binding affinity

One of the prominent features of IRE is the C1:G5 intra-loop base pairing (Fig. 2), making the apical loop a classic example of pseudotriloop [27, 28]. We have studied the importance of the C1:G5 base pairing for IRE-IRP binding in yeast by evaluating the binding affinity between the G5A mutant FTL1 (Fig. 2) and IRPs. The result indicated that disruption of this intra-loop base-pairing affects the binding of IRP1 as shown by the lower 3-AT concentration at which growth was still observed (Fig. 3). Interestingly, restoration of the pseudotriloop by the compensatory C1U mutation (mutant FTL2) significantly increased binding of IRP1 relative to FTL-wt IRE as indicated by sustained growth at 10 mM 3-AT (Fig. 3). Binding of IRP2 was less influenced by the pseudotriloop conformation as disruption (FTL1) and restoration (FTL2) of the intra-loop base pair slightly reduced and increased growth respectively, relative to FTL-wt (Fig. 3).

### Relevance of U6 and G3 in IRE pseudotriloop for IRP binding

Four additional mutants of FTL IRE were constructed to investigate the role of conserved bases within the CAGWGH motif for binding to IRP1 and 2, respectively (Fig. 4 and 5). Deletion of U6 (mutant FTL3) did not reduce but potentially increased binding affinity to IRP1 whereas replacing U6 with C (mutant FTL4) led to increased binding of IRP1 (Fig. 4). Substitution of G3 by U either with (mutant FTL6) or without (mutant FTL5) an additional U6G mutation merely resulted in slightly lower binding of IRP1 relative to FTL-wt IRE (Fig. 4). A similar trend was observed for IRP2 in binding mutants FTL3-6 (Fig. 5), showing increased binding to U6C mutant FTL4 but no change in binding affinity to the other mutants.

**Figure 4.**
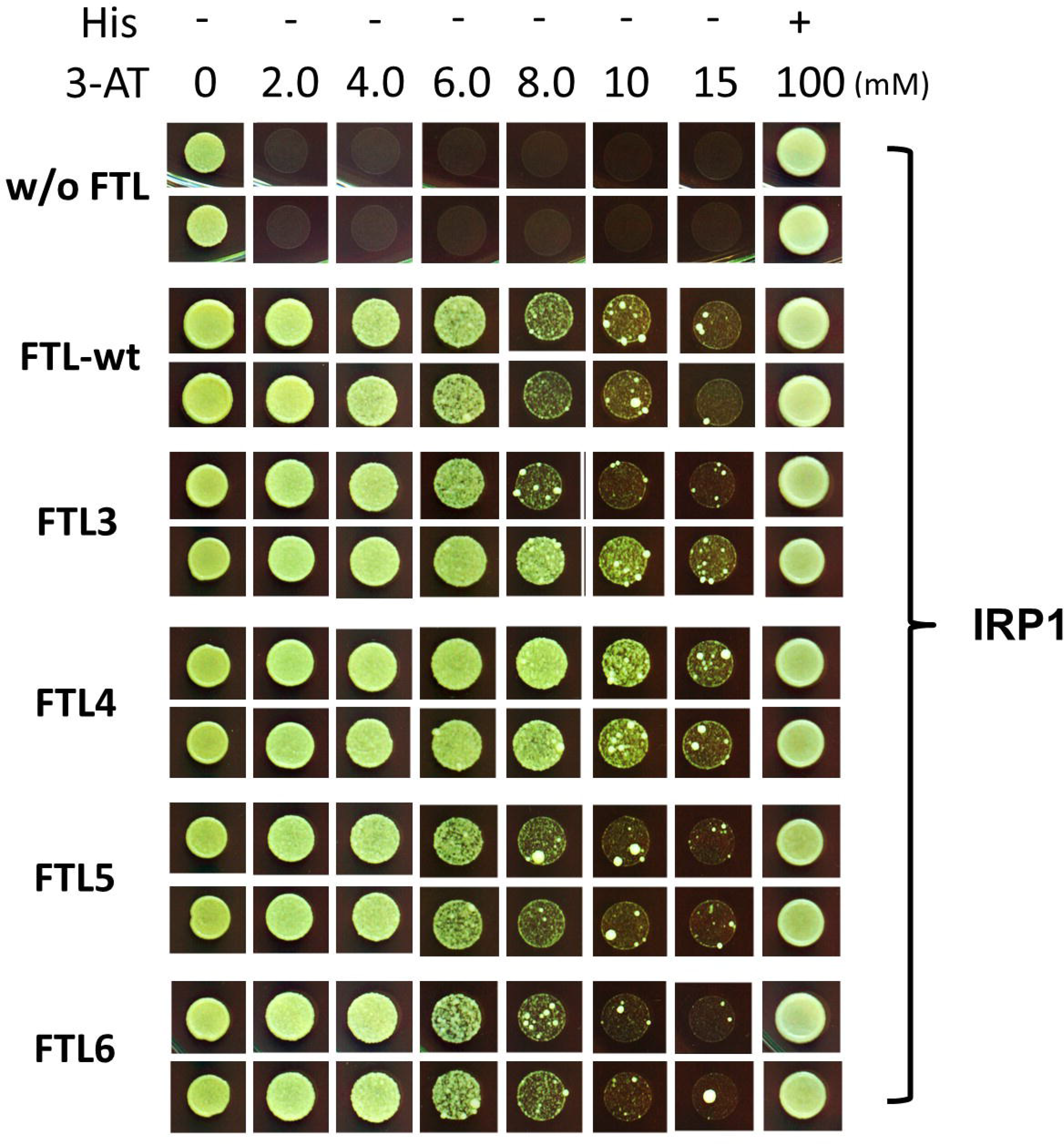
Yeast three-hybrid assay to verify the relevance of G3 and U6 of IRE pseudo-tri-loop to the binding with IRP1. The binding affinities between IRE mutants FTL3 to 6 and IRP1 were assayed by Y3H system, respectively. Clone w/o FTL is the negative control of which the RNA ligand has no IRE structure, while clones FTL3 to 6 contain IRE mutants shown in Fig. 2.

**Figure 5.**
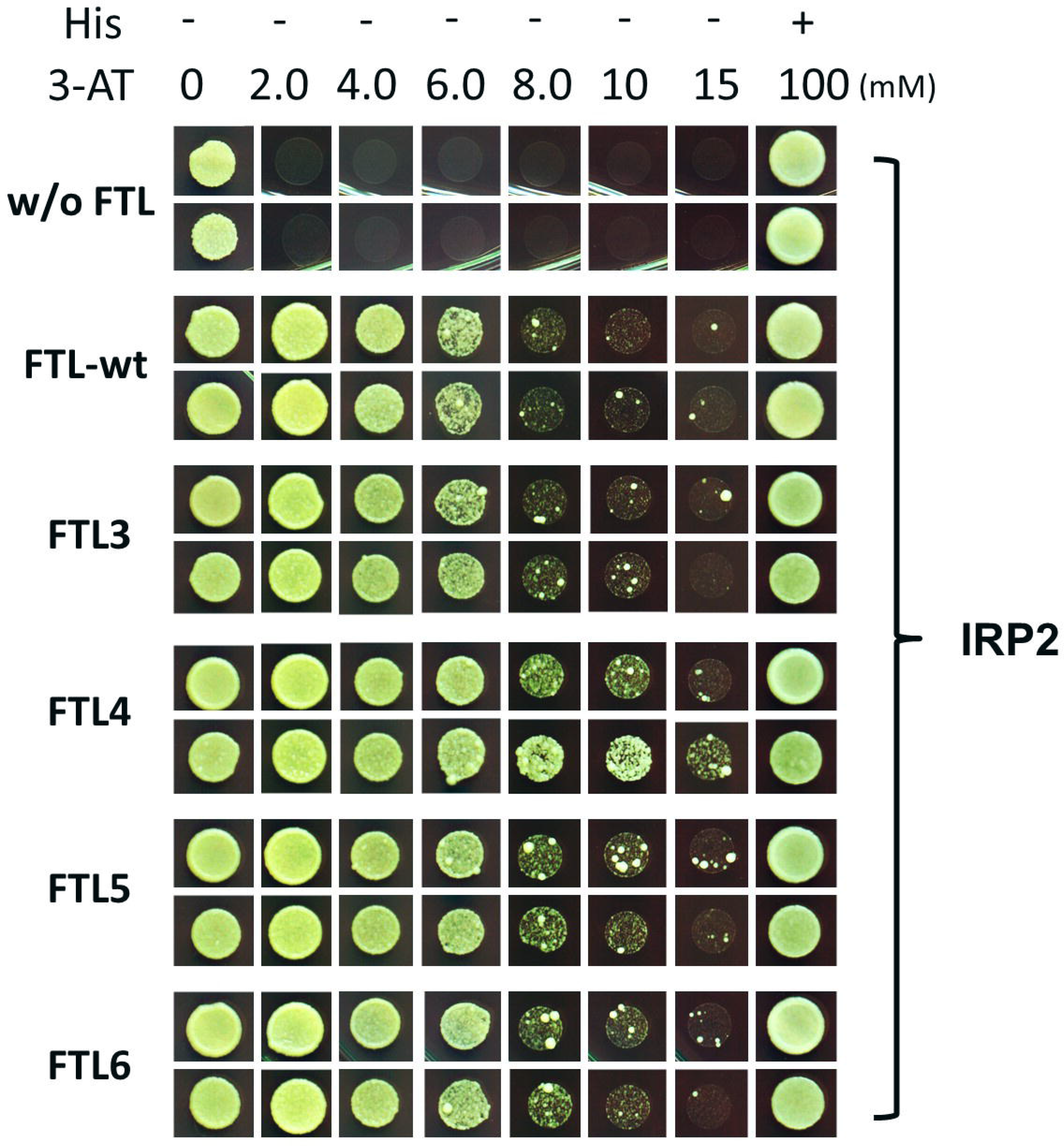
sYeast three-hybrid assay to verify the relevance of G3 and U6 of IRE pseudo-tri-loop to the binding with IRP2. The binding affinities between IRE mutants FTL3 to 6 and IRP2 were assayed by Y3H system, respectively. See legend to Fig. 3 for further details.

### Validation of the binding between IRE-like structures and IRPs

Several recently identified IRE-like structures (Fig. 6) were validated for their binding affinity to IRPs in Y3H system, including the IREs of EPAS1, α-Synuclein, SLC40A1, ALAS2, and SLC11A2 mRNA. All these RNA structures exhibit the characteristic features of an IRE. Figure 7 shows that IRP1 binds strongly to the variant IREs of SLC40A1, EPAS1, and ALAS2, weakly to the IRE of SLC11A2, and not to the proposed IRE of α-Synuclein. Compared to FTL-wt IRE, IRP1 binds to ALAS2 IRE with higher affinity, to SLC40A1 and EPAS1 IREs with similar affinity, and to SLC11A2 IRE with lower affinity. The stronger binding with ALAS2 is probably due to the bulge being a C, as mutant FTL4 (C-bulge) also has higher affinity than wild type IRE (Fig. 4). We conclude that IRE-like structures having a relatively stable 5-basepair upper stem interact with IRP1 with similar affinity as FTL IRE, while structures with a thermodynamically unstable upper stem, *e.g*. IREs of α-Synuclein and SLC11A2 mRNA, have weak or no detectable IRP1 binding in Y3H system. IRP2, however, binds to SLC11A2 IRE more strongly than IRP1 does, yet shows similar trends in binding to other variant IREs (Fig. 8).

**Figure 6.**
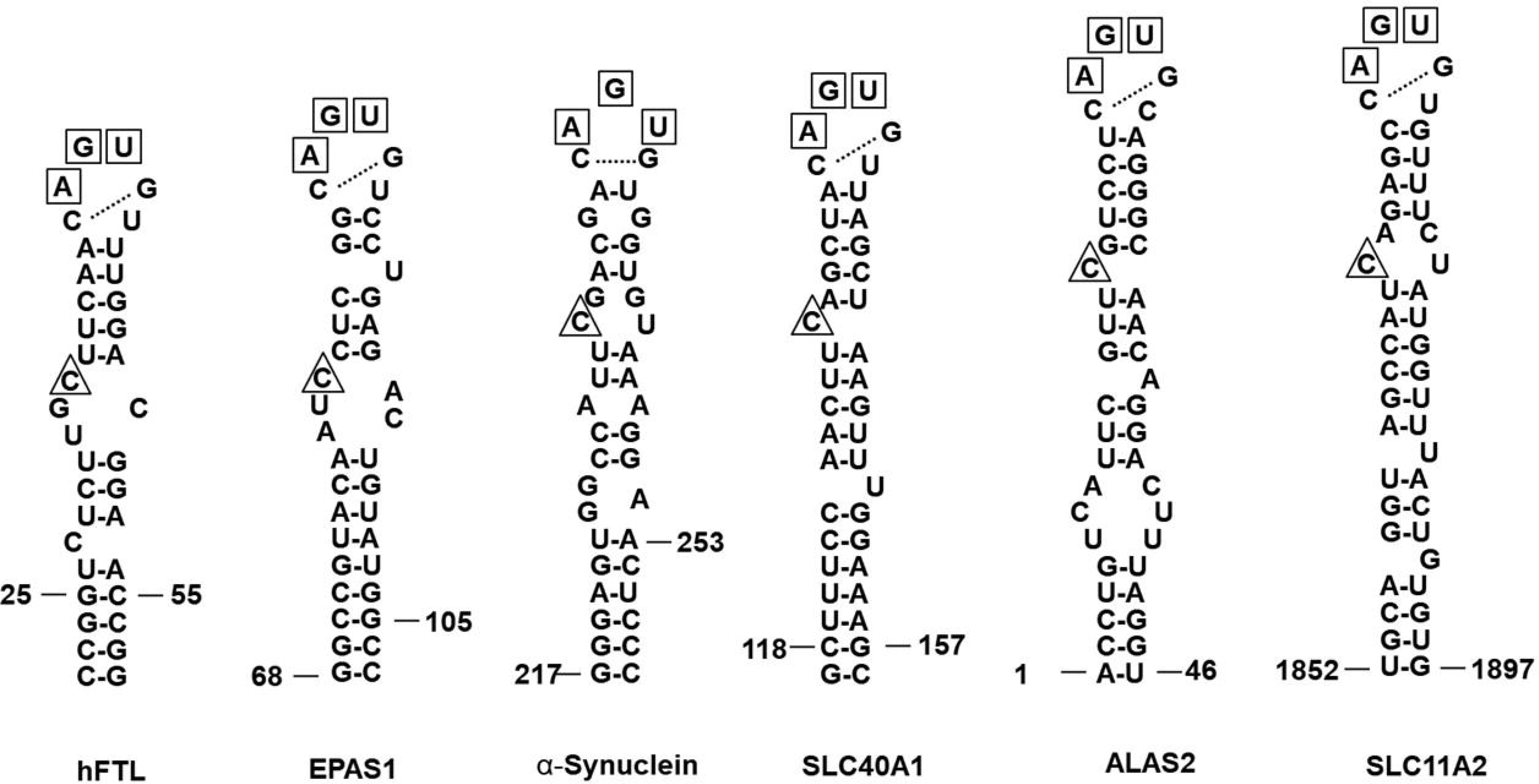
Variant IREs studied in yeast three-hybrid system. Secondary structures of Ferritin light chain (FTL) IRE [13], SLC40A1 IRE [39], Endothelial PAS domain protein 1 (EPAS1) IRE [15], α-Synuclein IRE [19], and 5′-aminolevulinate synthase 2 (ALAS2) IRE [40] are shown. The highly conserved A, G, and U bases in the IRE pseudotriloop are boxed, and the numbering indicates the span of each element according to the reference mRNA sequences of each gene (Accession number NM_000146.4, NM_001430.5, NM_000345.3, NM_014585.5, NM_000032.5, NM_000617.2. Additional nucleotides were introduced to the lower stem region as indicated to stabilize the IRE structure.

**Figure 7.**
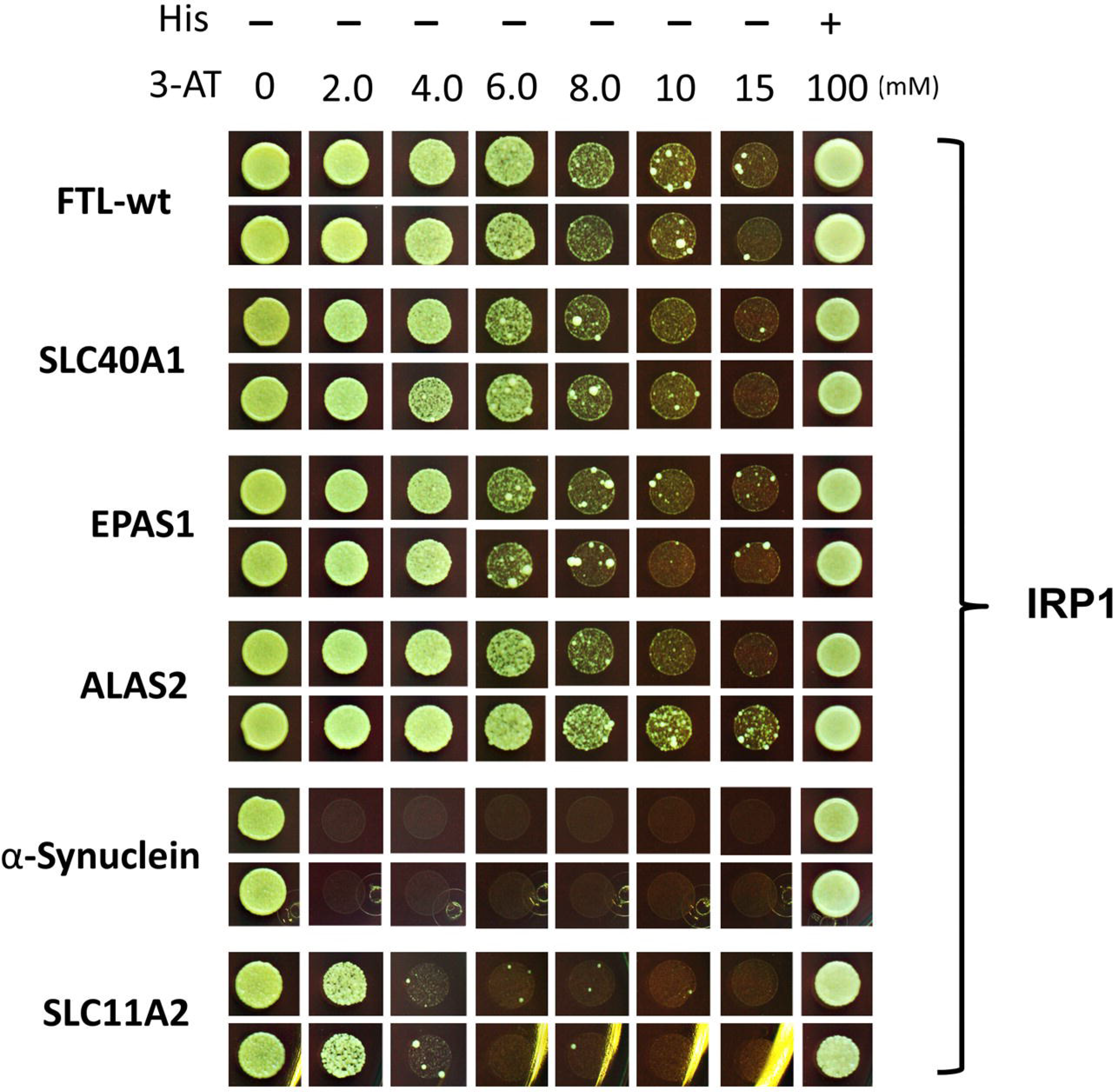
Yeast three-hybrid assay to verify the binding affinity between variant IREs and IRP1. Secondary structures of variant IREs are depicted in Fig. 6. See legend to Fig. 3 for further details.

**Figure 8.**
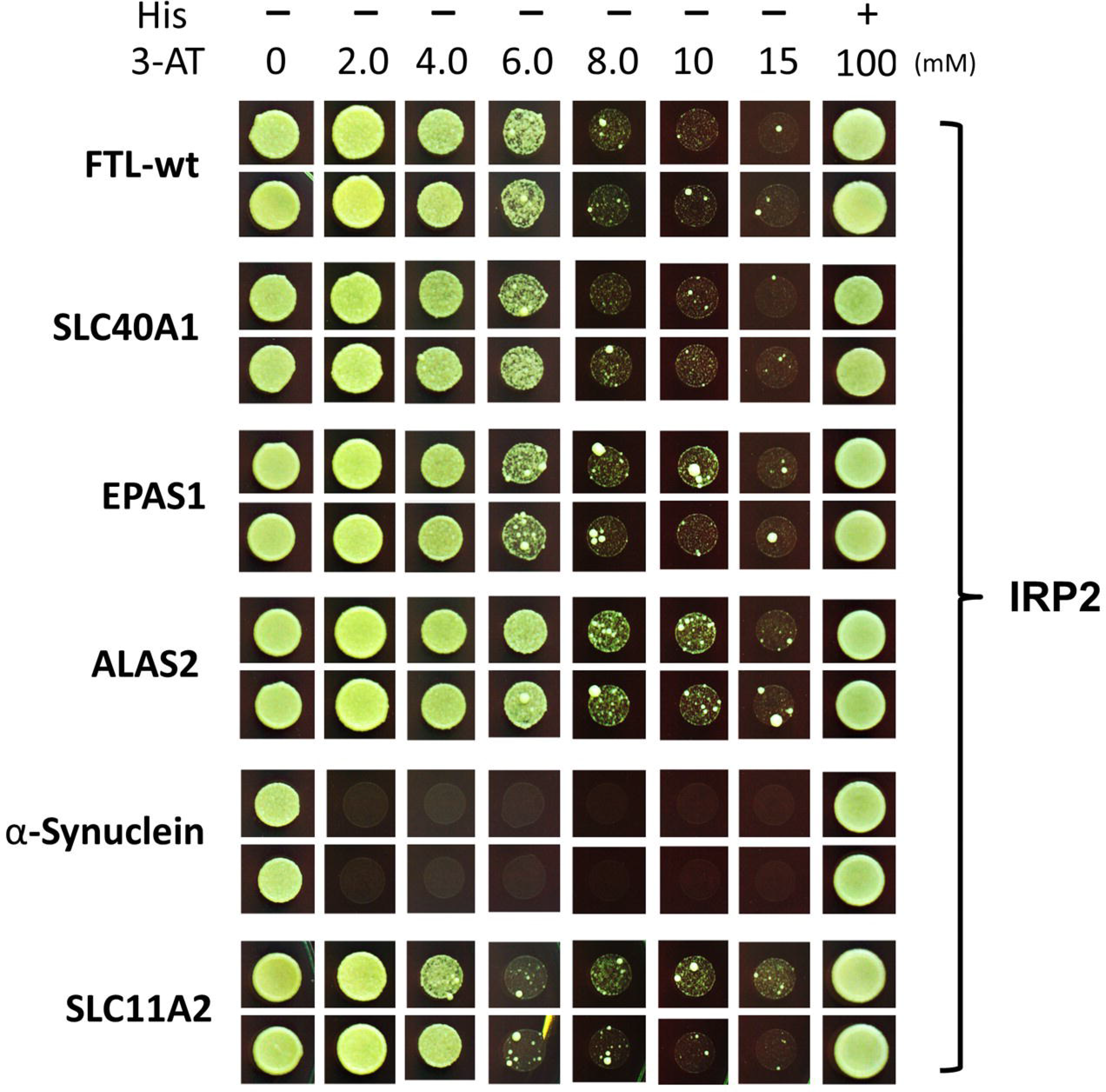
Yeast three-hybrid assay to verify the binding affinity between variant IREs and IRP2. Secondary structures of variant IREs are depicted in Fig. 6. See legend to Fig. 3 for further details.

## Discussion

### Insights in binding affinities between IRPs and variant IREs *in vivo*

In the studies of IRP-IRE complex reported previously, two large and spatially separate domains of IRP1 have been shown to bind to the IRE and predominantly form 10 bonds with the AGU pseudotriloop and 8 bonds with the inter-stem C-bulge [12, 29], suggesting that the pseudo-tri-loop and the C-bulge of IREs are the critical binding sites for IRP1. All the IRE variants studied in this study exhibit such a characteristic AGU pseudotriloop and the C-bulge (Fig. 6), but their binding affinities for IRPs were found to differ substantially in Y3H. For instance, EPAS1 and ALAS2 IREs were bound equally well by IRP1 as FTL IRE (Fig. 7), while the IREs of SCL11A2 and α-Synuclein showed less and no detectable binding to IRP1, respectively (Fig. 7). Recently, Khan *et al.* [30] reported that the kinetic and thermodynamic properties of ferritin and mitochondrial aconitase IRE-IRP1 interaction *in vitro* is affected by temperature and the presence of Mn^2+^. This discovery further suggested that *in vitro* and *in vivo* RNA-protein binding assays, *e.g*. EMSA and Y3H, may lead to different conclusions due to different assay conditions. Goforth *et al.*, [31] reported that IRP1 affinities for EPAS1 and ALAS2 IREs were 3.0 and 6.3 fold lower, respectively, than for the FTL IRE. However, we did not observe such binding differences *in vivo* (Fig. 7, 8), possibly because our Y3H assay is not sensitive enough to measure such changes. On the other hand, we found that the presence of the apical AGU sequence and the bulged C is not a guarantee for strong binding to IRPs; direct binding between the proposed IRE-like structure for α-Synuclein mRNA [7, 18, 19, 32], and IRPs was not detected in our assay (Fig. 7, 8). This result is in agreement with the previously reported inability to identify IRPs through RNA pulldown with the IRE-like structure of α-Synuclein and the inability to detect α-Synuclein mRNA by protein pulldown with IRP1/2 [16, 33]. These data suggest that the iron-mediated translational regulation of α-Synuclein mRNAs does not directly or solely rely on IRP-IRE interactions and may involve other proteins like eIF4F [34], or uses other regulatory mechanisms.

### Differential binding affinities between IRPs and variant IREs

Of the two IRP isoforms, IRP1 functions both as an RNA-binding protein and as the cytosolic aconitase, while IRP2 functions solely as an RNA-binding protein. It is generally believed that IRP1 is the major iron sensor and IRE binder as the abundance of IRP1 is higher than that of IRP2 in most cells [35]. However, IRP2 was also reported to be the dominant regulator of iron uptake and heme biosynthesis in erythroid progenitor cells by controlling the expression of TfR1 and ALAS2 mRNAs [7]. Recently, IRP1 and 2 were found to regulate, respectively and/or collaboratively, different cellular mechanisms of iron metabolism, tumorigenesis, and neurodegeneration [36–38]. In our Y3H system, we have found that the differences in IRP1 and 2 to bind the canonical IREs, *e.g*. FTL and SLC40A1 IREs, were insignificant. However, with respect to certain variant IREs and mutants, *e.g*. SLC11A2 IRE and the C1:G5 to U1:A5 mutant FTL2 tested in this study, IRP1 and 2 did show differential binding affinities. We hypothesize that such selectivity to the variant IREs *in vivo* potentially allows IRP1 and 2 to regulate different subsets of mRNA in cells, of which the IREs are preferentially bound by either IRP1 or IRP2. Further research is needed to prove this.

Altogether, our data show that Y3H can be used to reveal differences in the binding modes of IRP1 and 2, as well as to measure binding affinities of IRPs to IREs in a semi-quantitative manner and validate putative IRE-like structures.

